# Combining RADseq and contact zone analysis to decipher cryptic diversification in reptiles: insights from *Acanthodactylus erythrurus* (Reptilia: Lacertidae)

**DOI:** 10.1101/2022.09.30.510260

**Authors:** Paul Doniol-Valcroze, Loïs Rancilhac, José Carlos Brito, Aurélien Miralles, Philippe Geniez, Laure Benoit, Anne Loiseau, Raphaël Leblois, Christophe Dufresnes, Pierre-André Crochet

**Affiliations:** CEFE, CNRS, Univ Montpellier, EPHE, IRD, Montpellier, France; Zoological Institute, Technische Universität Braunschweig, 38106 Braunschweig, Germany; Animal Ecology, Department of Ecology and Genetics, Evolutionary Biology Centre, Uppsala University, Uppsala, Sweden; CIBIO, Centro de Investigação em Biodiversidade e Recursos Genéticos, InBIO Laboratório Associado, Campus de Vairão, Universidade do Porto, Vairão, Portugal; BIOPOLIS Program in Genomics, Biodiversity and Land Planning, CIBIO, Campus de Vairão, Vairão, Portugal; Department of Biology, Faculty of Sciences, University of Porto, Porto, Portugal; Institut de Systématique, Evolution, Biodiversité (ISYEB), Muséum national d’Histoire naturelle, CNRS, Sorbonne Université, EPHE, Université des Antilles, Paris, France; CEFE, EPHE-PSL, CNRS, Université de Montpellier, Biogéographie et Ecologie des Vertébrés, Montpellier, France; CBGP, INRAE, CIRAD, IRD, Montpellier SupAgro, University of Montpellier, Montpellier, France; LASER, College of Biology and the Environment, Nanjing Forestry University, Nanjing, People’s Republic of China

**Keywords:** *Acanthodactylus erythrurus*, Spiny-footed Lizard, Cryptic Speciation, Hybrid Zone, RADseq, Admixture

## Abstract

Linnaean and Wallacean shortfalls (Uncertainties on species taxonomy and distribution, respectively) are major factors hampering efficient conservation planning in the current context of biodiversity erosion. These shortfalls concern even widespread and abundant species in relatively well-studied regions such as the Mediterranean biodiversity hotspot which still hosts a large fraction of unrecognised biodiversity, notably in small vertebrates. Species delimitations have long been based on phylogenetic analyses of a small number of standard markers, but accurate lineage identification in this context can be obscured by incomplete lineage sorting, introgression or isolation by distance. Recently, integrative approaches coupling various sets of characters or analyses of contact zones aiming at estimating reproductive isolation (RI) have been advocated instead. Analyses of introgression patterns in contact zone with genomic data represent a powerful way to confirm the existence of independent lineages and estimate the strength of their RI at the same time. The Spiny-footed Lizard *Acanthodactylus erythrurus* (Schinz, 1833) is widespread in the Iberian Peninsula and the Maghreb and exhibits a large amount of genetic diversity, although the precise number and distribution of its genetic lineages remain poorly understood. We applied a RADseq approach to obtain a genome wide SNPs dataset on a contact zone in central Morocco between the previously described Rif and Middle-Atlas lineages. We show that these two lineages exhibit strong RI across this contact zone, as shown by the limited amount and restricted spatial extant of gene flow. We interpret these results as evidence for species-level divergence of these two lineages. Our study confirms the usefulness of RADseq approaches applied on contact zones for cryptic diversity studies and therefore to resolve Linnaean and Wallacean shortfalls.

## INTRODUCTION

Uncertainties on species taxonomy and distribution (Linnaean and Wallacean shortfalls, respectively, Lomolino 2004; Hortal et al. 2015) are major factors hampering efficient conservation planning in the current context of biodiversity erosion (e.g. Bini et al. 2006; Cardoso et al. 2011). Linnaean and Wallacean shortfalls do not only affect poorly studies organisms or remote biomes but are still widespread in conspicuous and abundant species in relatively well studied regions. For example, in the Mediterranean biodiversity hotspot, a large fraction of the species diversity of small terrestrial reptiles remains (or remained until recently) unrecognised (e.g. Kotsakiozi et al. 2018, Psonis et al. 2018, Kornilios et al. 2020, Kiourtsoglou et al. 2021), especially but not only in North Africa and in the Middle East (e.g. Kyriazi et al. 2008; Kapli et al. 2015; Miralles et al. 2020; Montgelard et al. 2020; Liz et al. 2021; Pizzigalli et al. 2021). Efficiently addressing Linnaean biodiversity shortfalls relies on using accurate methods to identify independent evolutionary lineages and assess their taxonomic ranks.

Phylogenetic analyses applied to standard molecular markers have significantly improved the accuracy and efficiency of species discovery and delimitation, but their results can be obscured by various processes such as incomplete lineage sorting, introgression and isolation by distance (IBD), especially when using limited genetic data (small number of loci and/or samples). Species delimitation models based on the multispecies coalescent have been proposed as an accurate way to objectively delimit species from multi-locus data, but they detect population structure rather than species limits (Sukumaran & Knowles 2017), making the use of complementary source of information necessary to assess the systematic ranks of the units identified by these models. In this context, two non-exclusive approaches can be used to resolve species boundaries: 1) using an integrative approach to assess the concordance and level of the divergences with other sets of characters (e.g. Miralles et al. 2013) and 2) assess reproductive isolation (RI) by quantifying admixture at secondary contact zones and determining how far introgression can be detected away from the core range of the lineages, allowing for a direct application of the biological species concept (and by extension the unified species concept, de Quieroz 2007, see Hillis 2019 for a discussion in herpetology).

Analyses of introgression patterns at contact zones, based on genome-wide markers, have recently been popularised to investigate species boundaries in many taxa (e.g. Shipham et al. 2019; Dufresnes & Martinez-Solano 2020, Dufresnes et al. 2020, 2021a, Lucek et al. 2020), including squamates (e.g. Schield et al. 2017; Caeiro-Dias et al. 2021a). The main factors limiting the applicability of this approach to poorly studied organism are the need for the existence of contact zones (i.e. it does not work on allopatric taxa), its reliance on comprehensive sampling schemes and on large multi-locus datasets. However, recent developments of genome reduction approaches and reductions of sequencing costs now allow the generation of genome-wide phylogeographic datasets even in non-model organisms. Restriction-sites Associated DNA sequencing (RADseq) and derived approaches have proven especially relevant for these studies, as they require little *a priori* genomic information, yield genotypes at many independent loci and at relatively low costs. Consequently, RADseq-based investigations of contact zones have been conducted in many systems to refine species boundaries (Caeiro-Dias et al. 2021a; b; Dufresnes et al. 2019; 2021a; Dufresnes & Martinez-Solano 2020). In this study, we use this approach to investigate a secondary contact zone of a Moroccan lizard, where previous multi-locus analyses suggested low levels of admixture (Rancilhac et al. 2022).

The Spiny-footed Lizard *Acanthodactylus erythrurus* (Schinz, 1833) is widespread in the Iberian Peninsula and the Maghreb, from Morocco to Tunisia (Fonseca et al. 2009), where it occupies a wide diversity of Mediterranean and semi-arid habitats with open grounds from the Atlantic coastal plains to the high Atlas Mountains (Miralles et al. 2020). Marked variations in morphology and ecology (from coastal sands to open forests through steppes) have led to the recognition of the coastal Moroccan populations as a distinct species (*A. lineomaculatus*) and of several subspecies within *A. erythrurus* (Bons & Geniez 1995). However, phylogenetic reconstructions based on mitochondrial DNA (mtDNA) did not support this taxonomy and suggested considerable cryptic diversity within the group (Harris et al. 2004; Fonseca et al. 2008; 2009; Beddek et al. 2018). More recently, the analyses of one mtDNA and nine nuclear (nDNA) markers by Miralles et al. (2020) highlighted the existence of five major phylogeographic lineages: an Ibero-Moroccan clade (Iberian Peninsula and most of Morocco), a Central Algerian clade, an Algero-Tunisian clade (Tunisia and coastal populations of Eastern Algeria, including populations described under the name *A. blanci*), and two clades from the Eastern and Western high Atlas newly described as *A. lacrymae* and *A. montanus* by these authors. The Ibero-Moroccan, Central Algerian and Algero-Tunisian lineages are deeply diverged and paraphyletic, granting further investigations of their taxonomic status, although they remain aggregated under the name *Acanthodactylus erythrurus*, including *A. lineomaculatus*. Furthermore, the Iberian-Moroccan clade is subdivided into 11 mitochondrial lineages (Miralles et al. 2020), although in depth analyses of nine nuclear loci brought only weak support to this phylogeographic pattern, which is obscured by high levels of alleles sharing (Rancilhac et al. 2022). Rancilhac et al. (2022) also demonstrated that at least part of the phylogeographic structure is imputable to Isolation by Distance (IBD), even if sampling gaps and low informativeness of the markers used precluded a reliable differentiation of IBD from vicariance. The delimitation of the evolutionary units within the Ibero-Moroccan *A. erythrurus* thus remains incomplete, hampering both our understanding of the species evolutionary history and taxonomic decisions to formally recognize this diversity.

Especially of interest, one contact zone between two highly divergent mitochondrial lineages has been identified near Ifrane, in the north of the Middle-Atlas (Miralles et al. 2020, Rancilhac et 2022). There, a lineage widely distributed across the Middle-Atlas (Fig. 1; subsequently referred to as Mid-Atl) occurs at close range from individuals belonging to a lineage occurring in the Rif Mountain range (subsequently referred to as Rif). This contact zone coincides with a steep transition in several morphological characters (Bons & Geniez 1995), including a fully diagnostic scalation feature (presence or absence of a small scale inserted between the subocular plate and the lip). Consequently, the two lineages have been placed into different subspecies *A. e. belli* (Rif) and *A. e. atlanticus* (Mid-Atl). They are characterized by deep mitochondrial divergences (c. 7.5 million years ago [Mya]), whereas nDNA analysis failed to recover them as reciprocally monophyletic (Rancilhac et al. 2022). Furthermore, statistical testing of IBD did not unambiguously support that these two groups are separated by a barrier to gene flow (Rancilhac et al. 2022). While these analyses are limited by the restricted dataset employed and sampling gaps, the contrast between mitochondrial and nuclear phylogeographic patterns is puzzling, and the existence of barriers to gene flow between the two groups remains to be confirmed. In case the barrier is confirmed, accurately quantifying admixture levels at the contact zone would allow to detect extrinsic barriers to gene flow (i.e. selection against hybrids) and determine whether the two lineages are different species.

**Figure 1.**
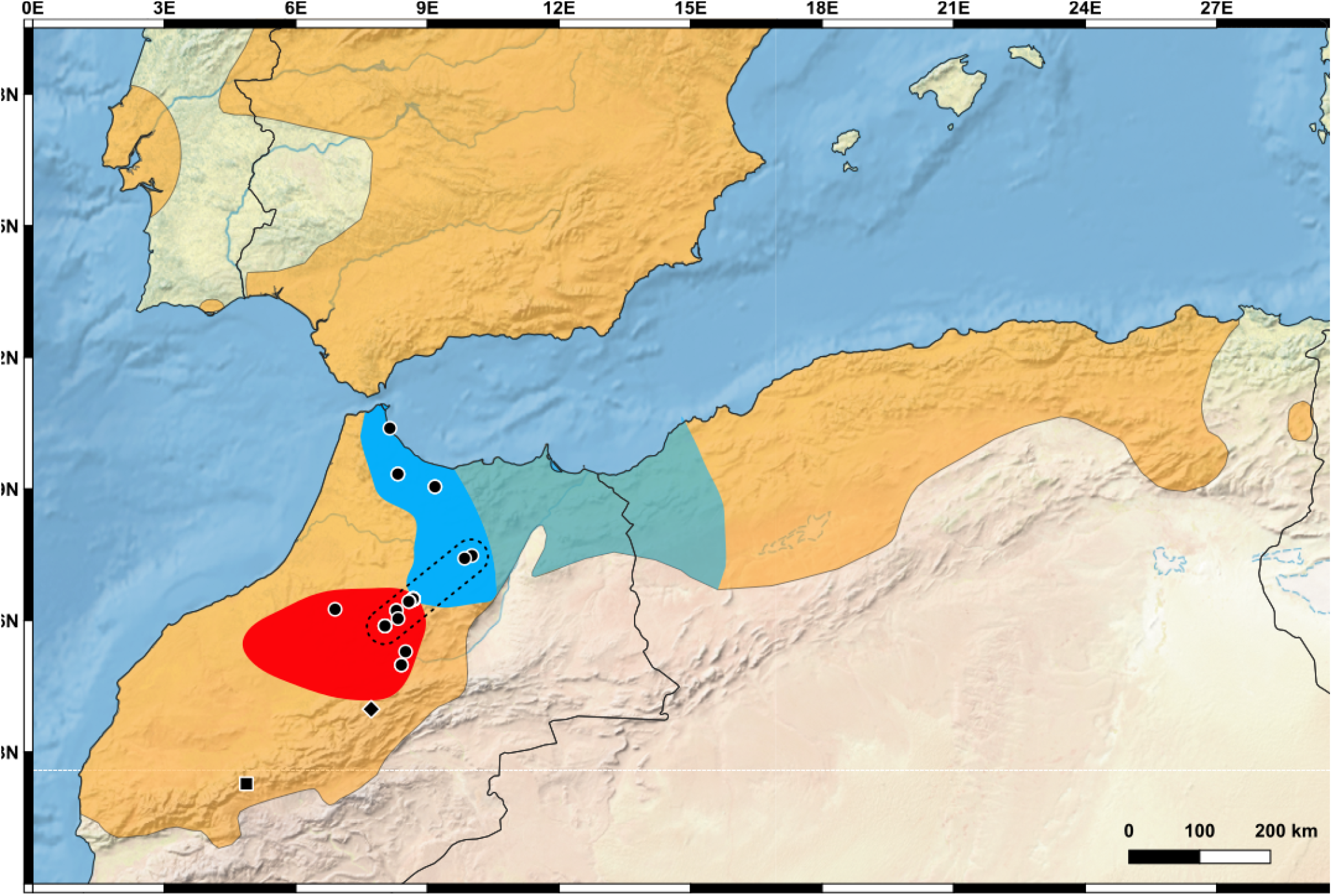
Distribution of the samples used for this study, with the dotted line showing the position of the “transect” populations across the contact zone. The other populations are “core” populations. The coloured areas denote the distribution range of *Acanthodactylus erythrurus* s.l. in orange (IUCN range polygon), the known range of the Middle-Atlas lineage in red (Miralles et al. 2020, Rancilhac et al. 2022) and the known range of the Rif lineage based on mitochondrial DNA haplotypes (light blue) and on nuclear DNA (dark blue; Beddek et al. 2018, Miralles et al. 2020, Rancilhac et al. 2022). The square and diamond symbols correspond to the outgroup *A.montanus* and *A. lacrymae*, respectively.

In this study, we generated genome-wide data through a RADseq approach from 35 individuals along the transition zone between the Rif and Mid-Atl lineages. Based on this data, we investigate overall population structure and genetic transition at the contact zone to clarify 1) whether the Middle-Atlas and Rif populations belong to vicariant lineages and 2) whether these lineages are isolated enough to deserve species status.

## MATERIAL & METHODS

### Sampling

A total of 35 individuals from the *Acanthodactylus erythrurus* complex, kept in the collection of the « Biogeography and Ecology of Vertebrates » team in Montpellier (CNRS & Ecole Pratique des Hautes Etudes, BEV-EPHE, UMR 5175-CEFE Montpellier) and already included in the study by Rancilhac et al. (2022), were selected for this study (Table 1). These individuals were sampled during dedicated field trips in Morocco between 2004 and 2011 at 16 separate locations (Fig. 1). Sampled locations cover two previously identified genetically and morphologically distinct lineages, Rif (Rif Massif and Northern Middle Atlas to Ifrah Lake) and Mid-Atl (Middle-Atlas south of the Ifrah Lake; Bons & Geniez 1995; Miralles et al. 2020). However, it should be noted that uncertainty remain as to whether the ranges of these lineages are wider than our sampling, since the identity of many other Moroccan populations remain unclear (Beddek et al. 2018, Miralles et al. 2020, Rancilhac et al. 2022). For each lineage, sampled localities are divided into two categories: localities in the core range (“core localities”, 80 – 200 km from the identified contact zone, five samples in Rif and Mid-Atl each), where individuals are likely less affected by introgression, and localities along a transect through the contact zone (“transect localities”, five samples from Rif and 20 from Mid-Atl) with a special sampling effort at the contact zone itself. The contact zone is located just north of the Ifrah Lake (Dayet Ifrah, approximately 15 km ENE of the town of Ifrane, in the Middle Atlas (see Figure 1). One individual of each of the recently described *A. lacrymae* and *A. montanus*, sampled in the Hight-Atlas were included as outgroups (Table 1, Fig. 1).

**Table 1:** Origin and geographic position of samples along the transect and probability of STRUCTURE assignment for K = 2. Coordinates are in decimal degrees (WGS84). Samples in bold are from the contact zone near Ifrah lake.

| Sample number | Individual identifier | Lineage | Latitude | Longitude | Likelihood of Rif assignment | Likelihood of MA assignment |
| --- | --- | --- | --- | --- | --- | --- |
| BEV.11873 | MART3 | RIF | 35.6379 | -5.2776 | 1 | 0 |
| BEV.11898 | KETA2 | RIF | 34.9373 | -4.6158 | 1 | 0 |
| BEV.11899 | KETA3 | RIF | 34.9373 | -4.6158 | 1 | 0 |
| BEV.11830 | TAZA1 | RIF | 35.0899 | -5.1595 | 1 | 0 |
| BEV.11831 | TAZA2 | RIF | 35.0899 | -5.1595 | 1 | 0 |
| BEV.T11837 | TZKA11 | RIF | 34.0708 | -4.1806 | 1 | 0 |
| BEV.T11836 | TZKA10 | RIF | 34.1042 | -4.0724 | 1 | 0 |
| BEV.11925 | **IFRA1** | RIF | 33.5828 | -4.9352 | 0.897 | 0.103 |
| BEV.11926 | **IFRA2** | RIF | 33.5828 | -4.9352 | 0.933 | 0.067 |
| BEV.11927 | **IFRA3** | RIF | 33.5828 | -4.9352 | 0.939 | 0.061 |
| BEV.11235 | OULM2 | MA | 33.4497 | -6.0812 | 0.001 | 0.999 |
| BEV.11238 | OULM5 | MA | 33.4497 | -6.0812 | 0 | 1 |
| BEV.11972 | BOUM4 | MA | 32.7655 | -5.1081 | 0 | 1 |
| BEV.11971 | BOUM5 | MA | 32.7655 | -5.1081 | 0 | 1 |
| BEV.11949 | CZAD1 | MA | 32.9297 | -5.0465 | 0 | 1 |
| BEV.11932 | **IFRA4** | MA | 33.5732 | -4.9315 | 0.012 | 0.988 |
| BEV.11933 | **IFRA5** | MA | 33.5732 | -4.9315 | 0.004 | 0.996 |
| BEV.11934 | **IFRA6** | MA | 33.5732 | -4.9315 | 0.016 | 0.984 |
| BEV.11935 | **IFRA7** | MA | 33.5732 | -4.9315 | 0.035 | 0.965 |
| BEV.11936 | **IFRA8** | MA | 33.5732 | -4.9315 | 0.015 | 0.985 |
| BEV.11937 | **IFRA9** | MA | 33.5732 | -4.9315 | 0.024 | 0.976 |
| BEV.11939 | **IFRA11** | MA | 33.5732 | -4.9315 | 0.022 | 0.978 |
| BEV.11940 | **IFRA12** | MA | 33.5732 | -4.9315 | 0.001 | 0.999 |
| BEV.11942 | **IFRA13** | MA | 33.5732 | -4.9315 | 0.024 | 0.976 |
| BEV.11944 | **IFRA14** | MA | 33.5732 | -4.9315 | 0.005 | 0.996 |
| BEV.11930 | **IFRA15** | MA | 33.577 | -4.9349 | 0.025 | 0.975 |
| BEV.11931 | **IFRA16** | MA | 33.577 | -4.9349 | 0.076 | 0.924 |
| BEV.T11809 | **IFRA17** | MA | 33.5449 | -5.0004 | 0.001 | 0.999 |
| BEV.11945 | **IFRA18** | MA | 33.5433 | -4.9927 | 0.001 | 0.999 |
| BEV.T11778 | AZRU4 | MA | 33.4351 | -5.1818 | 0 | 1 |
| BEV.T11779 | AZRU5 | MA | 33.4351 | -5.1818 | 0 | 1 |
| BEV.T11798 | AZRU21 | MA | 33.2458 | -5.3491 | 0.001 | 0.999 |
| BEV.T11800 | AZRU23 | MA | 33.2458 | -5.3491 | 0 | 1 |
| BEV.T11802 | AZRU25 | MA | 33.2458 | -5.3491 | 0 | 1 |
| BEV.T11797 | AZRU28 | MA | 33.3358 | -5.1570 | 0.001 | 0.999 |
| BEV.12001 | *A. lacrymae* | - | 32.2182 | -5.5501 | - | - |
| BEV.11794 | *A. montanus* | - | 31.2879 | -7.3824 | - | - |

### Molecular laboratory procedures

Total genomic DNA was extracted using the Dneasy Blood and Tissue Kit (QIAGEN, Hilden, Germany) following the manufacturer’s recommended procedures. RADseq libraries were prepared at the Centre de Biologie pour la Gestion des Populations (CBGP, Montpellier, France) following the methods described by Baird et al. (2008) and Etter et al. (2011), with some modifications as detailed below. For each sample, 180 ng of genomic DNA was digested with the frequent cutter restriction enzyme *SbfI*. P1 adapters (synthesised by IDT, USA) barcodes (Multiplex IDentifiers : MIDs) were then ligated to the digested fragments. Low sample diversity is an intrinsic problem for restriction enzyme based libraries (Krueger et al. 2011). To increase diversity within the library and reduce the number of phasing errors, which can be responsible for reduced sequence quality (Elshire et al. 2011), a set of 32 MIDs of 6 and 7 bp was used. The set was constructed to maximise the balance between nucleotides at each position and the MIDs differed by at least three nucleotides in order to reduce misassignments due to sequencing errors. Ligated products were pooled in four different pools and sheared using an Ultrasonicator SS220 (Covaris, Inc.; 10% duty cycle, intensity equal to 5, cycle/burst equal to 200, duration of 70 seconds) and fragments of 300-600 bp were size-selected and purified with AMPure XP beads (Agencourt), for concentration and removal of small fragments (e.g. unligated adaptors). After end-repair and 3’ A-tailing, P2 adapters (including Illumina primers for pair-end sequencing and 4 differents 6-7 nucleotides barcodes (MIDs)) were ligated. Final enrichment PCR amplification was performed in 5 independent 25 μL wells. The initial amount of DNA was reduced to 30 ng and the number of PCR cycles was reduced to 17 to limit the amount of PCR duplicates. PCR products were pooled and homogenised, and the obtained library was quantified by qPCR using the Library Quantification Kit-Illumina/Universal (KAPA; KK4824) for sequencing preparation. It was then denatured using NaOH and diluted to 7 pM prior to clustering. Clustering and 100nt pair-end read sequencing were performed on a single lane of a HiSeq 3000 flowcell at Toulouse GeT (Genotoul, Toulouse, France) following the manufacturer’s instructions. A low-concentration spike-in (1%) of PhiX Control v3 was used as an in-lane positive control for alignment calculations and quantification efficiency. Image analyses and base calling were performed using the HiSeq Control Software (HCS) and Real-Time Analysis (RTA) software (Illumina).

### RADseq data processing and assembly

All raw reads were demultiplexed and trimmed to 80 bp using *process_radtags* in Stacks (Catchen et al. 2013), after quality checking with FastQC (https://www.bioinformatics.babraham.ac.uk/projects/fastqc/). As *de novo* assembly of Pair-Ended RADseq data is challenging due to variable insert sizes (Etter et al. 2011), d*e novo* assembly of the RAD loci was only performed from the R1 reads, in Ipyrad v.0.9.50 (Eaton & Overcast 2020). Single-end sequences assembling does not allow PCR duplicates filtering, potentially impacting downstream issues (Andrews et al. 2014, 2016), but recent evidence suggest that this issue may not be as limiting as often thought (Euclide et al. 2020). The minimum depth for base calling was set to 8, and the optimal clustering threshold (CT, the minimal percentage of similarity for two sequences to be considered orthologous) was determined empirically to reduce the risk of introducing paralogs in the dataset by using an inappropriate value. To do so, the assembly was performed with CTs ranging from 0.85 to 0.99, and we observed the effect of varying the CT on 1) the number of recovered loci shared across ≥80 % of the samples, 2) the number of Parsimony Informative Sites (PIS) in the assembly and 3) the proportion of missing data in the assembly. Both intra-samples CT (i.e. threshold used to cluster reads within samples) and between-samples CT (i.e. threshold used to cluster loci across individuals) were treated as the same value, as we are working at the intraspecific level, where we do not expect alleles from separate populations to be much more divergent than alleles within individuals. Ultimately, we selected a CT value that would maximize the number of PIS and loci shared by ≥80 % of the samples, while minimizing the proportion of missing data. The other assembly parameters were left to default.

Two assemblies were performed: 1) Assembly 1 includes all 35 *A. erythrurus* individuals plus one sample each of *A. montanus* and *A. lacrymae* as outgroups for phylogenetic inference; 2) Assembly 2 does not include outgroups in order to mitigate the effects of loci dropout, and all loci with missing individuals were filtered out for population genetics analyses. Both assemblies were formatted to obtain complete loci sequences and unlinked phased genotypes corresponding to the SNP with highest coverage at each locus (or randomly drawn in case of equality).

### Phylogenetic reconstruction

The concatenation matrix from Assembly 1 was used to perform maximum likelihood (ML) phylogenetic inference. The analyses were conducted in IQTREE v. 1.6.8 (Nguyen et al. 2015) under a GTR+Γ substitution model, and branch support was assessed using the RH approximate likelihood test (aLRT) with 1,000 pseudoreplicates. The obtained topology was rooted using *A. montanus* and *A. lacrymae* as hierarchical outgroups.

### Population structure and contact zone analyses

The phylogeographic structure of our focal species was further investigated in two ways. First, the relative genetic distances among individuals were inferred using a Principal Component Analysis (PCA) computed on SNPs genotypes. This analysis was performed with the *adegenet 2*.*1*.*7* R package (Jombart 2008) based on the unlinked SNPs from Assembly 2. Secondly, the Bayesian clustering algorithm STRUCTURE 2.3.4 (Pritchard et al. 2000) was used to infer the number of genetic groups in the data, and individual ancestries in these groups. Analyses were run with the number of groups (K) from 1 to 5, with four replicates for each. Most of the parameters were set to default values as advised by STRUTURE’s manual (Pritchard et al. 2000). We applied the admixture model, the correlated allele frequencies option and allowed the degree of admixture alpha to be inferred from the data as recommended by Falush et al. (2003). For each run, 10,000 burn-in steps and 100,000 iterations were performed. We used Evanno et al. (2005) method implemented in STRUCTURE HARVESTER (Earl & von Holdt 2012) to select the K value which maximised the DeltaK.

In order to quantify the geographic transition between the Rif and Mid-Atl lineages, we fitted sigmoid clines to assignment probabilities (Q) inferred by STRUCTURE, as well as to the allele frequencies of diagnostic SNPs (i.e., differentialy fixed in the “core populations” of the two lineages, see Fig. 1) in “transect populations”, using the R package *hzar 0*.*2-5* (Derryberry et al. 2014). Allele frequency clines are defined by their width (*w*) and center (*c*) and can be complemented by exponential tails to model long-range introgression (“stepped clines”). We fitted clines with width *w* and center *c* parameters for each diagnostic SNP and the Akaike Information Criterion (AIC) was computed from the likelihood estimates to select the one most appropriate to the hybrid zone.Cline parameters result from a balance between selection against hybrids (which reduces cline width) and dispersal (which increases cline width) and can thus be informative of the processes mediating patterns of genetic introgression between hybridizing species at their secondary contact zone (Barton 1979, Szymura & Barton 1986). For instance, narrow clines flanked by large introgression tails can reflect strong selection against hybrids yet with the diffusion of neutral alleles far away from the contact zone (e.g. Dufresnes et al. 2021a). Asymmetric clines, such as stronger introgression towards one species than the other, can be signatures of hybrid zone movements (Wielstra 2019).

## RESULTS

### RADseq data assembly and phylogenetic inference

After empirical optimization, the final assemblies were performed with a Clustering Threshold set to 95 % (Sup Mat 1) The Assembly 1 (with outgroups and missing data included) yielded 97,765 loci (totalling 7,353,579 bp), containing 477,106 SNPs of which 87,828 are considered independent (one per locus). The Assembly 2 (no outgroups, no missing data) included 13,395 loci (totalling 1,007,162 bp), containing 69,542 SNPs of which 13,007 are considered independent. The ML phylogenetic inference of Assembly 1 yielded a robust tree (Fig. 2), fully supporting the reciprocal monophyly of the Rif and Middle-Atlas populations.

**Figure 2.**
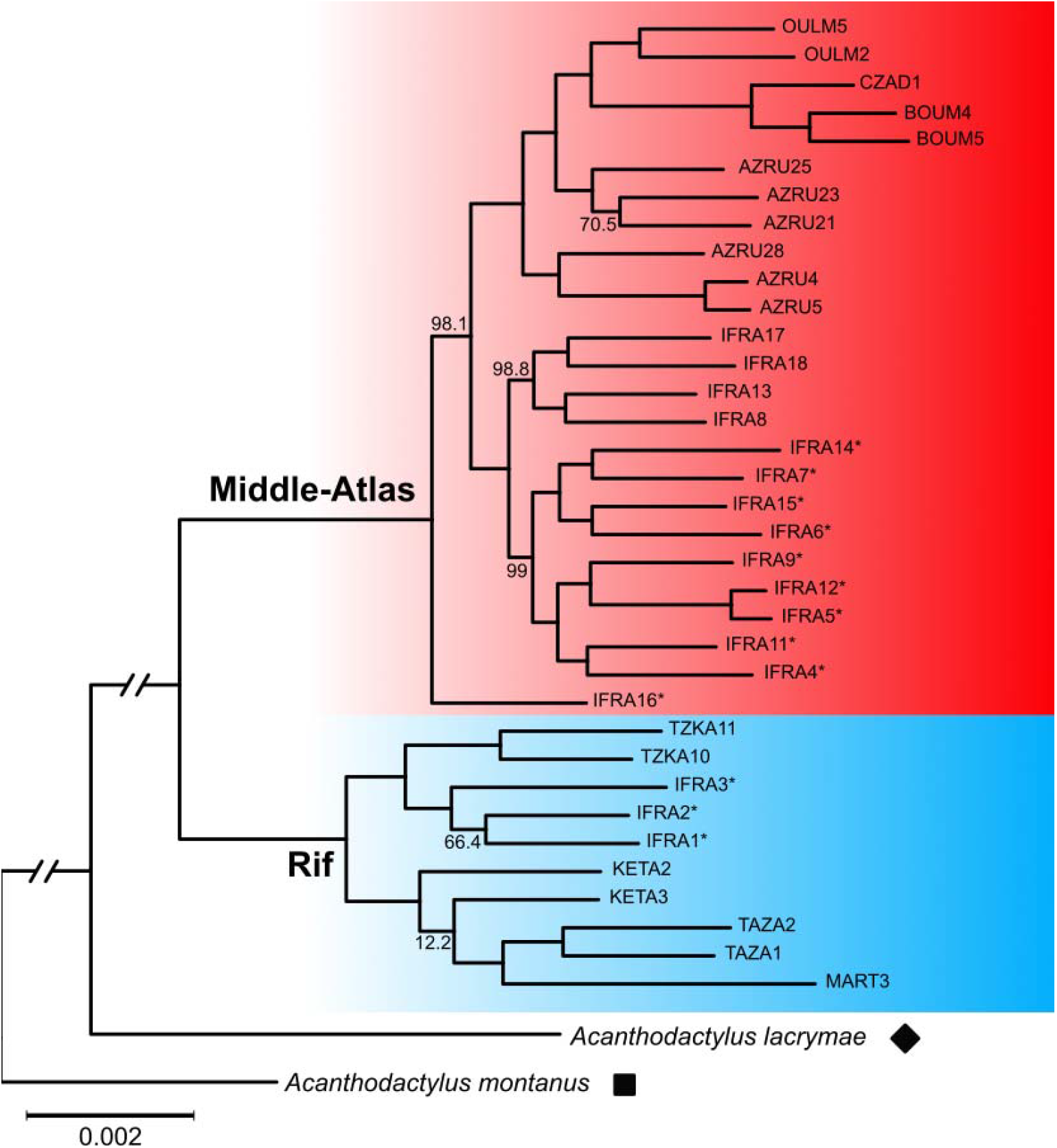
Maximum likelihood phylogenetic tree inferred from the concatenated RAD loci (7,353,579 bp). Approximate Likelihood ratio test (aLRT) supports are shown at the nodes when <100. Samples originating from the contact zone are indicated by asterisks.

### Analysis of the Rif / Middle-Atlas contact zone

The first two axes of the PCA explain respectively 11.24% and 5.10% of the total genetic variation. The Rif and Mid-Atl populations are clustered separately by PC1. PC2 shows additional structure within the Rif lineage, distinguishing individuals from the northern and southern parts (Fig. 3, corresponding to the Rif-N and Rif-S mitochondrial lineages in Rancilhac et al. 2022). In both lineages, some individuals from Ifrane (near the contact zone) show a somewhat intermediate position on PC1, consistent with low but detectable levels of introgression.

**Figure 3.**
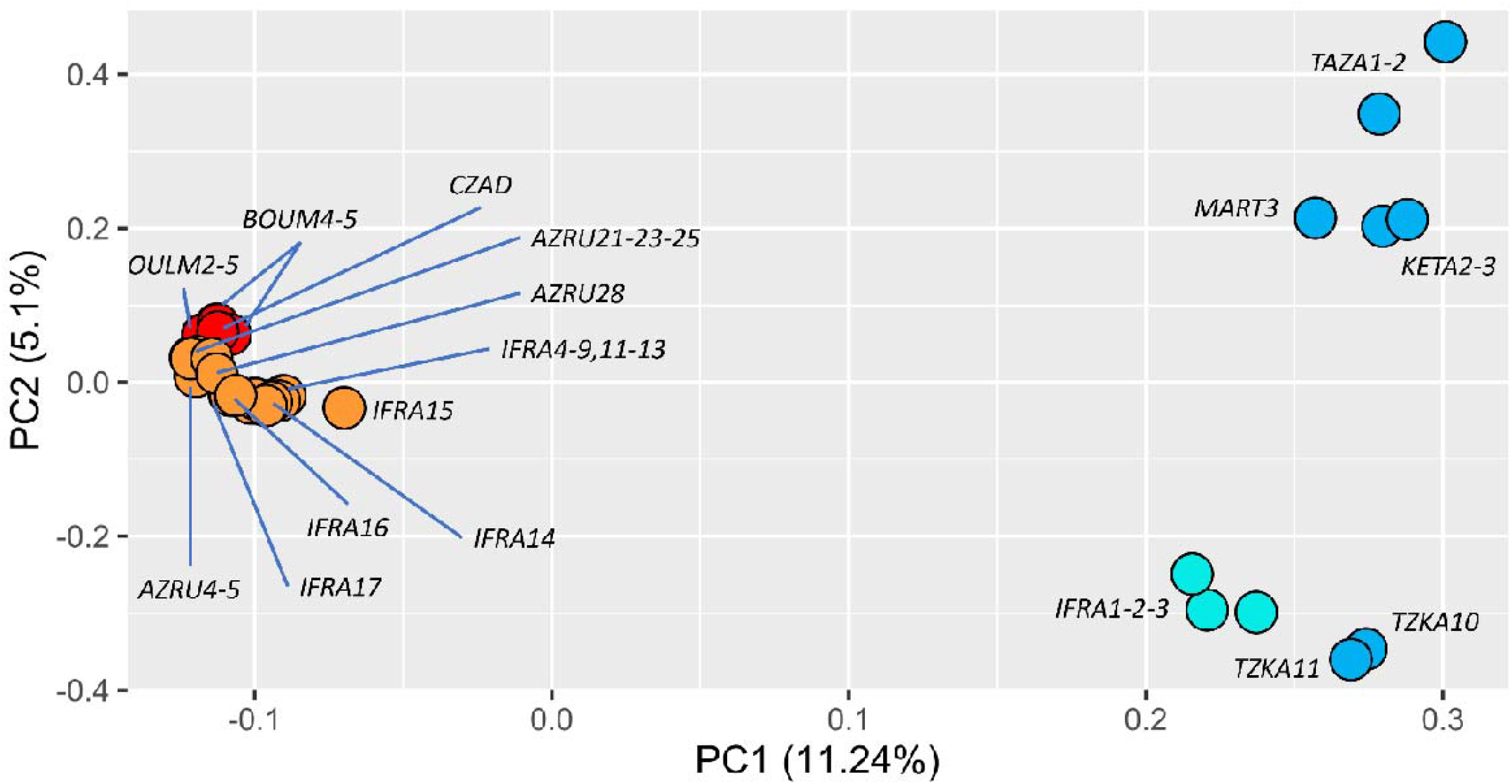
PCA performed on the independent SNPs matrix of Assembly 2. Red/orange and blue/light blue circles represent individuals sampled from “core” and “contact zone” populations of the Mid-Atl and Rif lineages respectively.

STRUCTURE analyses identified K = 2 as the best solution (Fig. 4A), corresponding to the Rif and Mid-Atl lineages. In this clustering solution, all samples had an assignment probability > 0.95 to either group (Fig. 4B, 4C, 4D), except for several samples from the contact zone, which showed variable levels of genetic admixture (Table 1, Fig. 4B, 4C, 4D). In Ifrane, the three northernmost individuals (IFRA 1-2-3) were assigned to the Rif lineage with weak introgression from Mid-Atl (Q = 0.897, 0.933 and 0.939 respectively) while the 14 remaining individuals clustered with the Mid-Atl lineage and were either pure (Q > 0.95, IFRA4, 5, 6, 7, 8, 9, 11, 12, 13, 14, 15, 17, 18) or weakly introgressed (IFRA16 ; Table 1, Fig. 4B).

**Figure 4.**
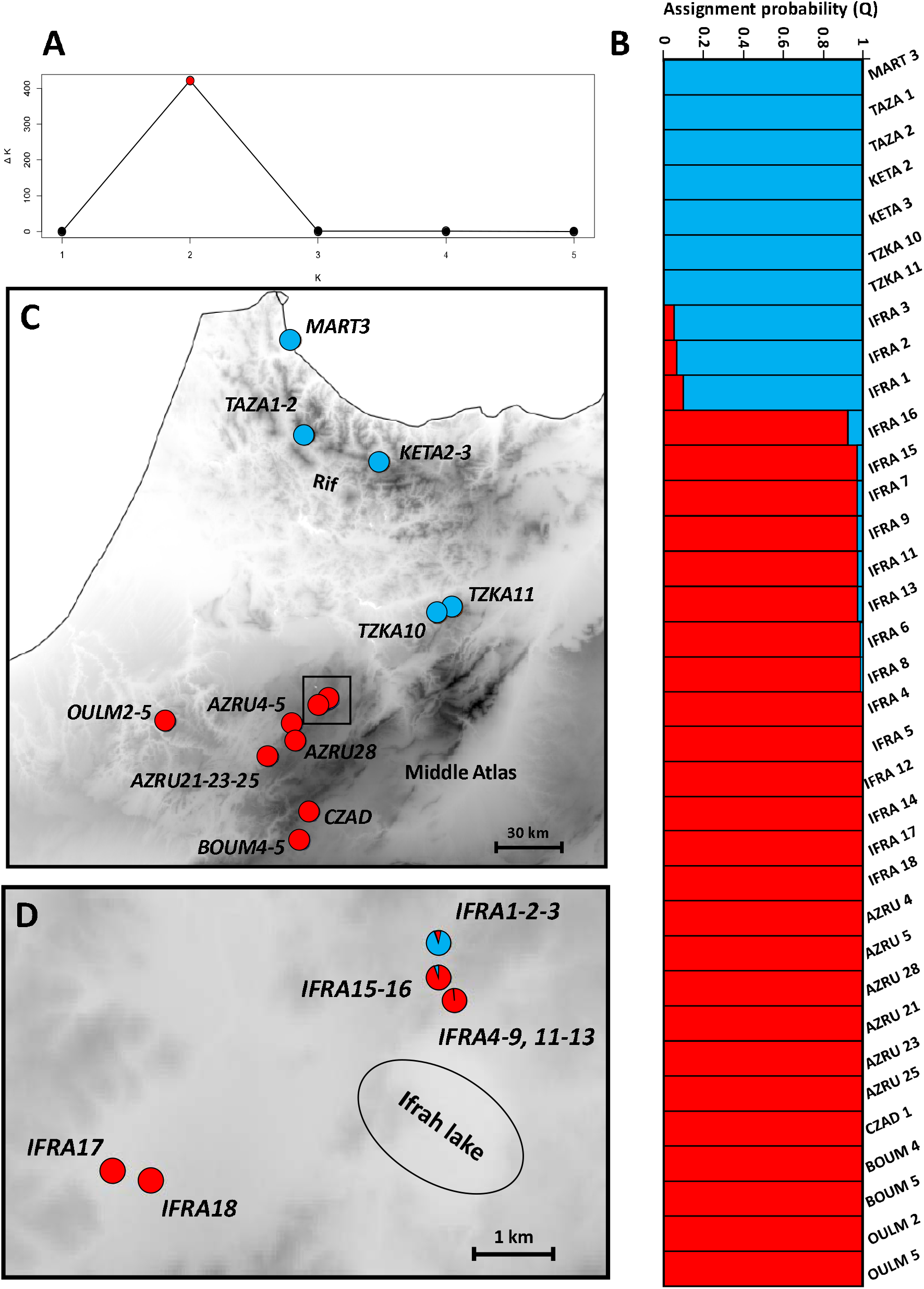
A) Variation of DeltaK for STRUCTURE runs with K ranging 1 – 5, calculated as described by Evanno et al. (2005). B) Assignment probabilities to the Rif (blue) and Mid-Atl (red) lineages for all individual, obtained from a STRUCTURE run with K = 2. Each bar represents an individual. C) Map of the study area showing the position of the sampling sites. The pie charts represent the average assignment probabilities of the sampled populations to each lineage (Rif in blue and Middle Atlas in red) obtained from a STRUCTURE analysis with K=2. D) Zoom on the contact zone around Ifrah Lake corresponding to the black square on A.

The best sigmoid cline model fitted on the average genomic ancestry (STRUCTURE’s Q) for the transect populations did not include introgression tails. This cline modelled a very sharp transition between the Rif and Mid-Atl lineages, with a width estimated to 1,260 m, and the centre located to the Ifrah Lake (Fig. 5A). Locus-by-locus clines, fitted to 1,820 diagnostic SNPs with (alleles frequencies of 0 or 1 in the core populations) mostly followed the genome average estimates, with 95% of widths spanning 600-42,000 m (median: 7,300 m), roughly following a Poisson distribution (Fig. 5 B-C). Most clines were asymmetric, with their centres shifted on the Rif side of the contact (negative distance positions on Fig. 5B), where we lacked a proper transect sampling (Fig. 4). Accordingly, 95% of the cline centres were located -19.4 – 2.5 km from the median centre.

**Figure 5.**
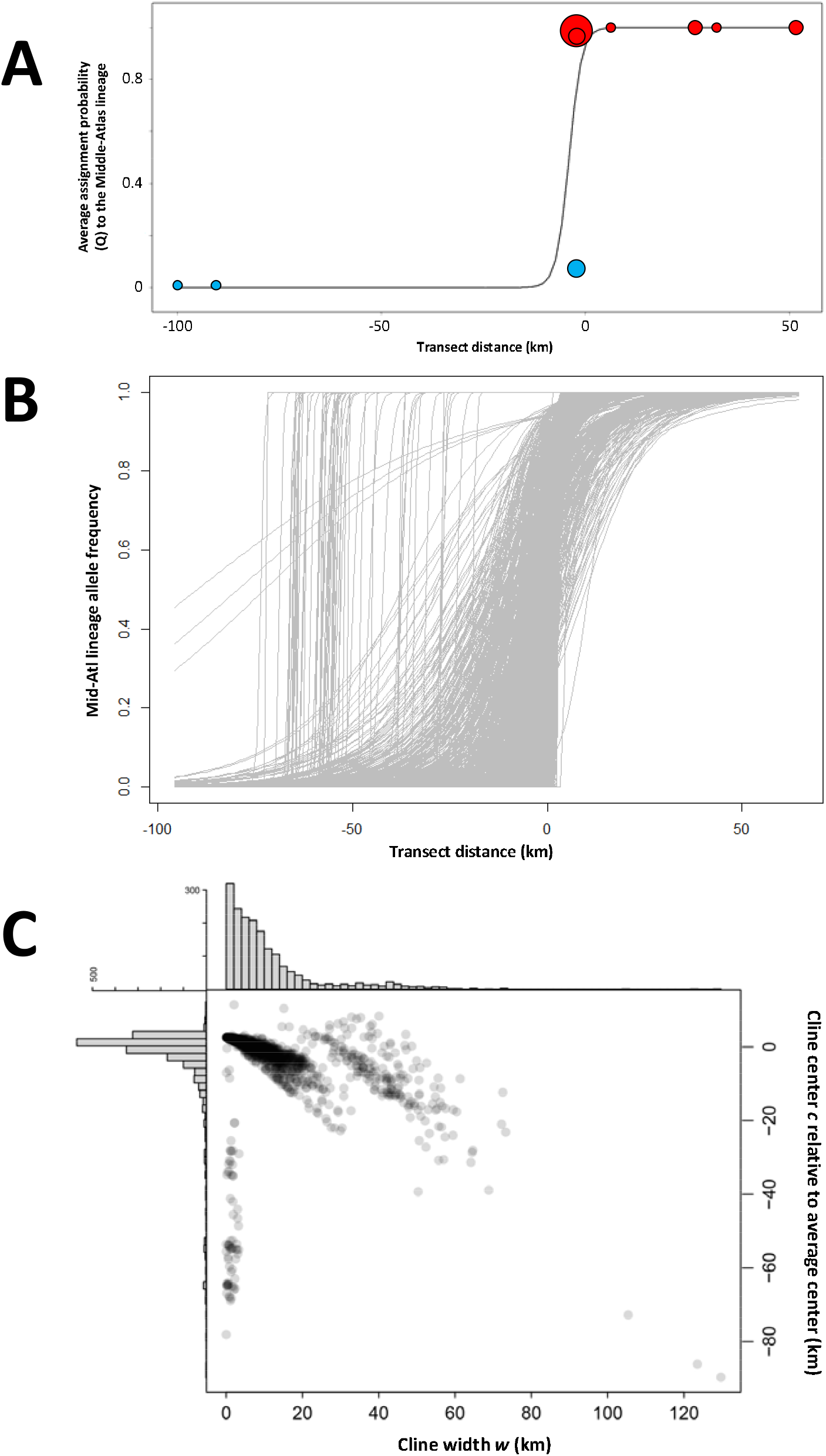
Geographic clines fitted across the Rif/Mid-Atl hybrid zone, including 10 populations along a broad north-south transect using the *hzar* package from: A) Genomic average as the average Q assignment probabilities from a STRUCTURE run with K=2. Red circles = the Mid-Atl lineage, blue circles = Rif lineage, their diameters indicating the number of individuals per population. B) Individual clines for 1820 diagnostic SNPs exhibiting alternative fixed alleles in core populations. C) Width *w* and centre *c* parameters for the individual clines.

## DISCUSSION

### *Genomics confirm a secondary contact between two species-level lineages within Moroccan populations of the* Acanthodactylus erythrurus *complex*

Previous studies using multi-locus data already highlighted the presence of several divergent phylogeographic lineages of *Acanthodactylus erythrurus* in Morocco, although the exact number of lineages involved has proven hard to tackle so far (Miralles et al. 2020, Rancilhac et al. 2022). Among these, the existence of a lineage inhabiting northern Morocco (here the Rif lineage) was only moderately supported by tests of vicariance in Rancilhac et al. (2022), although this Rif lineage exhibits a diagnostic scalation feature leading to its formal recognition as the subspecies *A. e. belli* in Bons and Geniez (1995). However, high levels of allele sharing made it difficult to estimate the levels of differentiation and RI between this lineage and the lineage inhabiting the Middle-Atlas and possibly the Atlantic coast of Morocco (Rancilhac et al. 2022). Here, using thousands of genome-wide markers, we confirmed that the “Rif” and “Mid-Atl” lineages form two distinct clusters without intermediate individuals away from their contact zones, as expected with vicariant units and confirming their independent evolution. Within the contact zone, individuals from both lineages were sampled as close as 600 m appart and although introgressive hybridization occurs, as revealed by the presence of admixed individuals, it occurs only at restricted geographic and genomic scales, consistently with substantial selection against hybrids. This pattern is most likely explained by an allopatric speciation scenario, where both lineages evolved separately long enough to accumulate mutations, ultimately leading to reproductive incompatibilities (Turelli et al. 2001, Pinho et al. 2009, Caeiro-Dias et al. 2021a, b). An alternative explanation would be that the secondary contact is very recent, so that even without reproductive isolation, introgression could still be unconspicuous across the 600m that currently separate the two lineages in our sample. This seems very unlikely given that no recent landscape changes that could have triggered a range expansion were recorded recently in the area and that *A. erythrurus* populations, already exhibiting the morphological features of the two lineages, have been known in the Dayet Ifrah for at least 30 years (PG pers. obs.). Strong ecological isolation also seems unlikely, as individuals of both lineages along the contact zone were found in the same, homogenous habitats of clear oak forest with extensive grassy clearings and edges.

The level of divergence and reproductive isolation between the Rif and Middle-Atlas populations, as indicated by the strong genome-wide population structure and low levels of introgression, respectively, is consistent with species rank under the biological species concept (and by extension the unified species concept, de Quieroz 1998, Mayr 2000, de Quieroz 2007), leading us to consider that the Ibero-Moroccan clade comprises at least two distinct species. This implies that *A. erythrurus* represents a species complex, as first illustrated by the discovery and description of *A. lacrymae* and *A. montanus* from the High Atlas by Miralles et al. (2020). The Ibero-Moroccan populations of *A. erythrurus* are further divided into several genetic groups with divergence levels seemingly comparable to that between the Mid-Atl and Rif lineages (Rancilhac et al. 2022). The generation of a large genome wide dataset covering an extensive sampling across the Maghreb is needed to accurately delimit species within the complex and determine their distributions. Although species delimitation will be more challenging for some of these lineages, given an anticipated absence of contact zones because of their patchy distributions, analysing morphological and molecular data alongside should allow a complete taxonomic revision of the complex. In anticipation of such global revision, and because uncertainy on the overall distributions of the Mid-Atl and Rif species make it difficult to thoroughly apply nomenclatural rules, we refrain from formally naming them here.

### RADseq analyses of secondary contact zones as a tool for species delimitation in non-model organisms

Our study supports the relevance of genome wide data in hybrid zones to infer species boundaries, even when previous multi-locus studies failed to unambiguously resolve them (here, Miralles et al. 2020, Rancilhac et al. 2022). Most of the geographic clines fitted to the genome hybrid index and to thousands of species-diagnostic SNPs all displayed a very steep transition. Moreover, the distribution of locus-specific width estimates roughly followed a Poisson distribution abutting zero, indicating that barrier loci, e.g. loci potentially involved in post-zygotic isolation, are scattered through the genome, a pattern characteristic of species boundaries in amphibians’ hybrid zones (Dufresnes et al. 2021a).

However, it should be noted that many clines appeared asymmetric, with their centres being shifted towards the Rif side of the transect (Fig. 5B, C). Shifts of cline centres can reflect a biological situation, namely differential introgression at specific loci caused by variation in selection parameters along loci (Barton 1979, Szymura & Barton 1986), but also by hybrid zone movement (Wielstra 2019) or drift (Jofre and Rosenthal 2021). It might also reflect unequal sampling along the transect: contrary to our dense and even sampling of Mid-Atl, a 100 km gap exists between the contact zone and the closest “core” Rif populations (TZKA 10 & 11). Precision of parameter estimations in cline analyses is sensitive to sampling gaps, so part of the variation in cline centre positions on the Rif side, where we have no sample close to the contact zone, might result from such inflation of confidence intervals. Some species-diagnostic loci could truly introgress further away than our sampled “contact zone” populations, but not as far as the reference populations, due to genuine differential introgression. This would result in cline centres shifted north more than their real positions because the precision of cline centres estimates north of the contact zone is limited by our sparse sampling. In addition, some of the markers considered as species diagnostic – because they featured allele frequency differences of 1 between our reference samples (located 100km away from the contact zone for the Rif lineage) – may in fact reflect intraspecific structure between the Rif edge populations of the contact and the Rif reference populations. This explanation would account for the fact that several outlier loci showed cline centres located halfway between these populations, e.g., ∼50km from the actual centre (Fig. 5B, C). Another limitation of our study is that we only used diagnostic loci to model cline parameters. This affects the inference of variation in introgression levels among loci as loci that introgress more than the genomic average are also more likely to introgress further away from the contact zone, resulting in shared alleles in core populations that will exclude them from the set of diagnostic loci. In other words, using only diagnostic loci underestimate the amount and geographic extent of genomic introgression and historical gene flow across the contact zone. However, it does not affect the inference of current patterns of reproductive isolation, which is based on genotypic composition of individuals within the contact zones.

Note that these limitations do not affect the conclusions of our study: most of the genome behaved like the genome average, and even if it remains unclear whether some loci truly introgress further away from the contact, this would not jeopardize the specific status of the Rif and Mid-Atl lineages. Even between mostly incompatible genomes, some neutral loci are expected to diffuse far within species ranges once recombination has broken down linkage between them and the barrier loci (Barton 1979, Polechová & Barton 2011). Accordingly, speciation is often achieved before interspecies gene flow entirely stops, so sharp clines may be paralleled by far-ranging introgression without questioning the taxonomic status of lineages (as shown in *Bombina* toads, Dufresnes et al. 2021b). However, the potential unreliability of the outlier clines reported here and the use of diagnostic loci only prevents the interpretation of the genomic landscape of introgression. We thus call for the careful design of hybrid zone studies, such as the inclusion of large numbers of reference specimens from different populations (in order to increase the chance to only flag lineage-diagnostic loci in analyses), dense yet even sampling along geographic transects (in order to improve the reliability of cline estimates) and inclusion of non-diagnostic loci if the genomic landscape if introgression is to be interpreted.

## Supporting information

Sup Mat 1

## ACKNOWLEDGEMENTS

Research conducted in the scope of the LIA “Biodiversity and Evolution”. Part of this work was carried out by using the resources of the national INRAe MIGALE (Migale bioinformatics Facility, doi: 10.15454/1.5572390655343293E12) and GENOTOUL (Bioinfo Genotoul, https://doi.org/10.15454/1.5572369328961167E12) bioinformatics HPC platforms, as well as the local Montpellier Bioinformatics Biodiversity (MBB, supported by the LabEx CeMEB ANR-10-LABX-04-01) and CBGP HPC platform services. RL was supported by the Agence Nationale de la Recherche (GENOSPACE ANR-16-CE02-0008 and INTROSPEC ANR-19-CE02-0011), and by recurrent funding from INRAE.

