## Supplementary material for "Combining RADseq and contact zone analysis to decipher cryptic diversification in reptiles: insights from *Acanthodactylus erythrurus* (Reptilia: Lacertidae)": Sup Mat 1

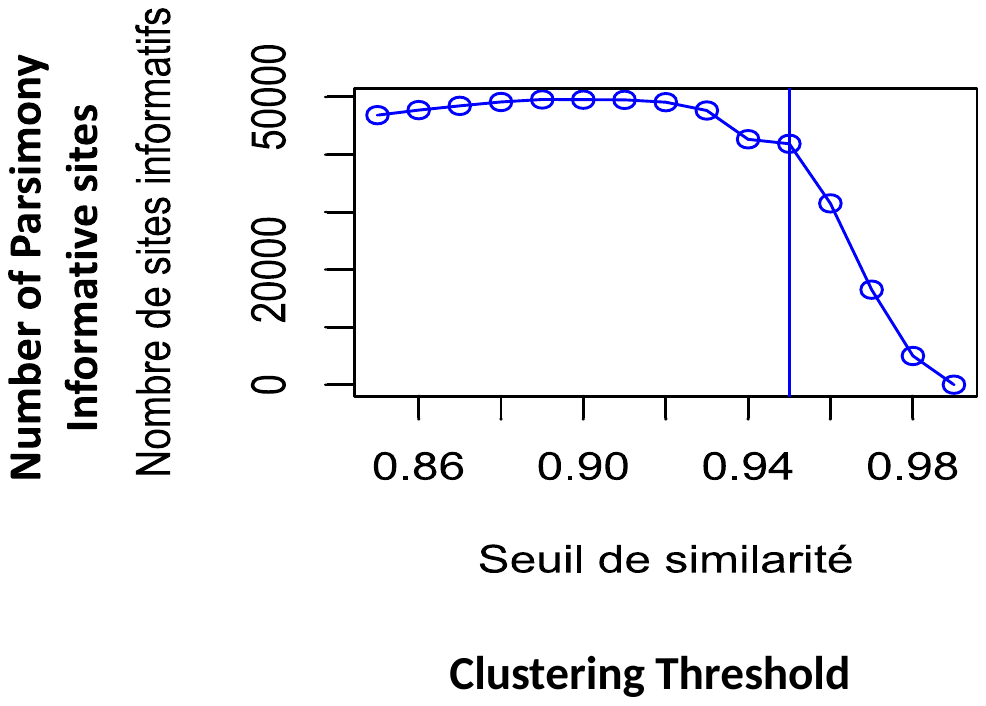

**Supplementary Material 1.** Graphical representation of the empirical optimisation of the clustering threshold. The chosen value (0.95) maximises the number of Parsimony Informative Sites without allowing a to low clustering threshold value to avoid the introduction of paralogs in the dataset.
